## Supplementary figures for "Nucleus accumbens D1- and D2-expressing neurons control the balance between feeding and activity-mediated energy expenditure"

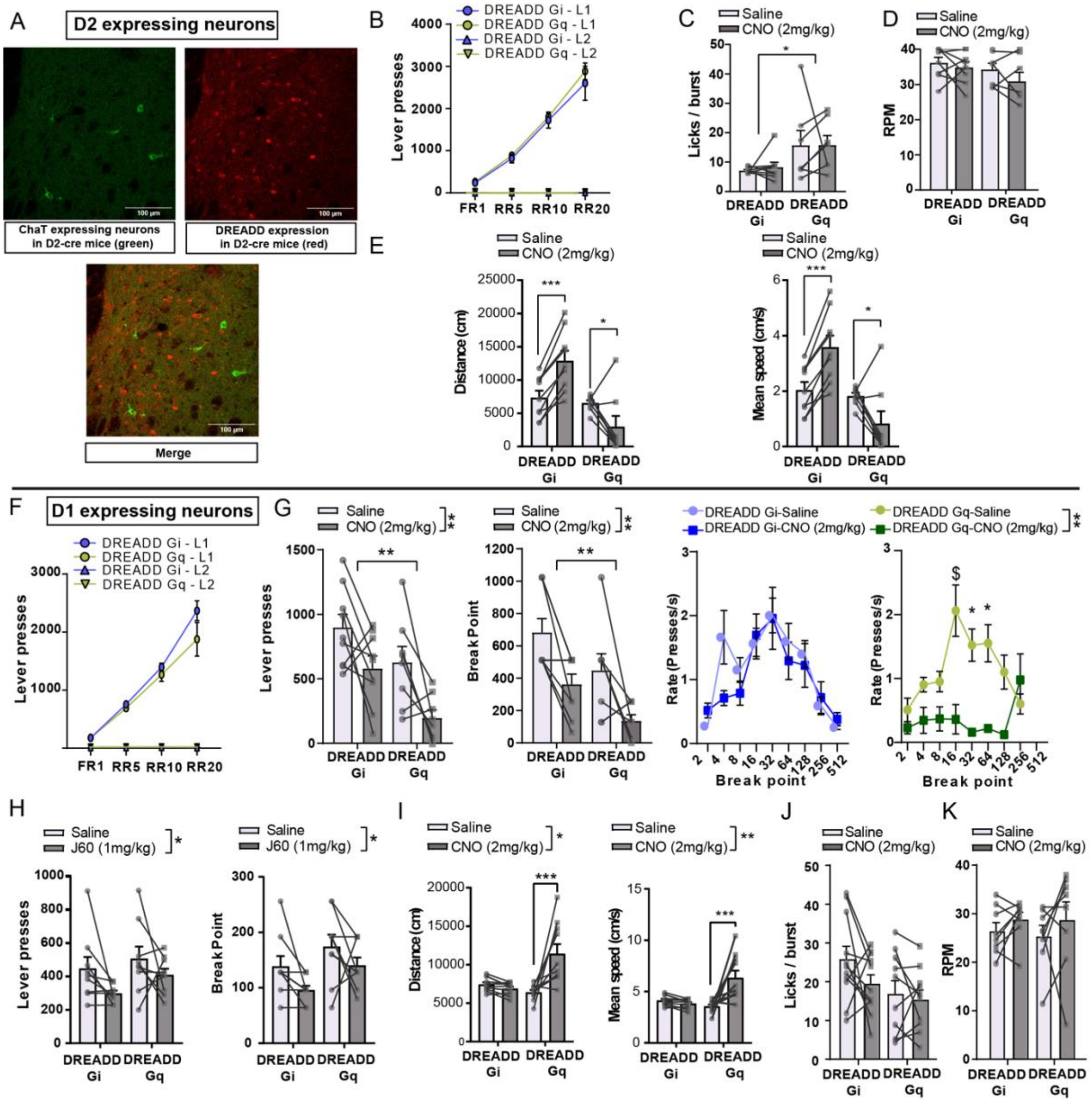

**Suppl. Figure 1 (relates to Figure 1):**

(A) Representative picture of NAc coronal section with immuno-labeled ChAT (Choline acetyl transferase) in green, (Top left), immuno-labeled DREADD-mCherry in red (Top right) and merge (Bottom). (B) Number of lever presses on the reinforced (L1) and non-reinforced (L2) levers across the different schedules of reinforcement in D2-cre mice expressing either DREADD Gi (n=9) or DREADD Gq (n=7) in the NAc, in the absence of CNO administration. (C) Licking microstructures (number of licks per burst) during consumption of a palatable solution (milk) in D2-cre mice expressing DREADD Gi (n=8) or Gq (n=7). (D) Rotation per min (RPM) in a rotarod in D2-cre mice under inhibition (DREADD Gi, n=8) or activation (DREADD Gq, n=7) of D2-neurons. (E) Distance (left) and mean speed (right) in an open field in D2-cre mice expressing DREADD Gi (n=9) or Gq (n=8). (F) Number of lever presses on the reinforced (L1) and non-reinforced (L2) levers across the different schedules of reinforcement in D1-cre mice expressing either DREADD Gi (n=9) or DREADD Gq (n=8) in the NAc, in the absence of CNO administration. (G) Effect of 2 mg/kg CNO in D1-cre mice expressing DREADDs Gi (n=9) or Gq (n=8) on the number of lever presses (left), breakpoint (middle) and ratio requirement (right) in the PR task. (H) Effect of the DREADD ligand JHU37160 (J60) in D1-cre mice expressing DREADDs Gi (n=9) or Gq (n=9) on the number of lever presses (left) and breakpoint (right) in the PR task. (I) Distance (left) and mean speed (right) in an open field in D1-cre mice expressing DREADD Gi (n=11) or Gq (n=10). (J) Licking microstructures (number of licks per burst) during consumption of a palatable solution (milk) under inhibition (DREADD Gi, n=11) or activation (DREADD Gq, n=10) of D1-neurons. (K) Rotation per min (RPM) in a rotarod in D1-cre mice under inhibition (DREADD Gi, n=8) or activation (DREADD Gq, n=8) of D2-neurons. \$: $0.05<p<0.1$ ; * : $p<0.05$ ; ** : $p<0.01$ ; *** : $p<0.001$ ; Error bars = s.e.m. See Table 2 for statistical analyses.

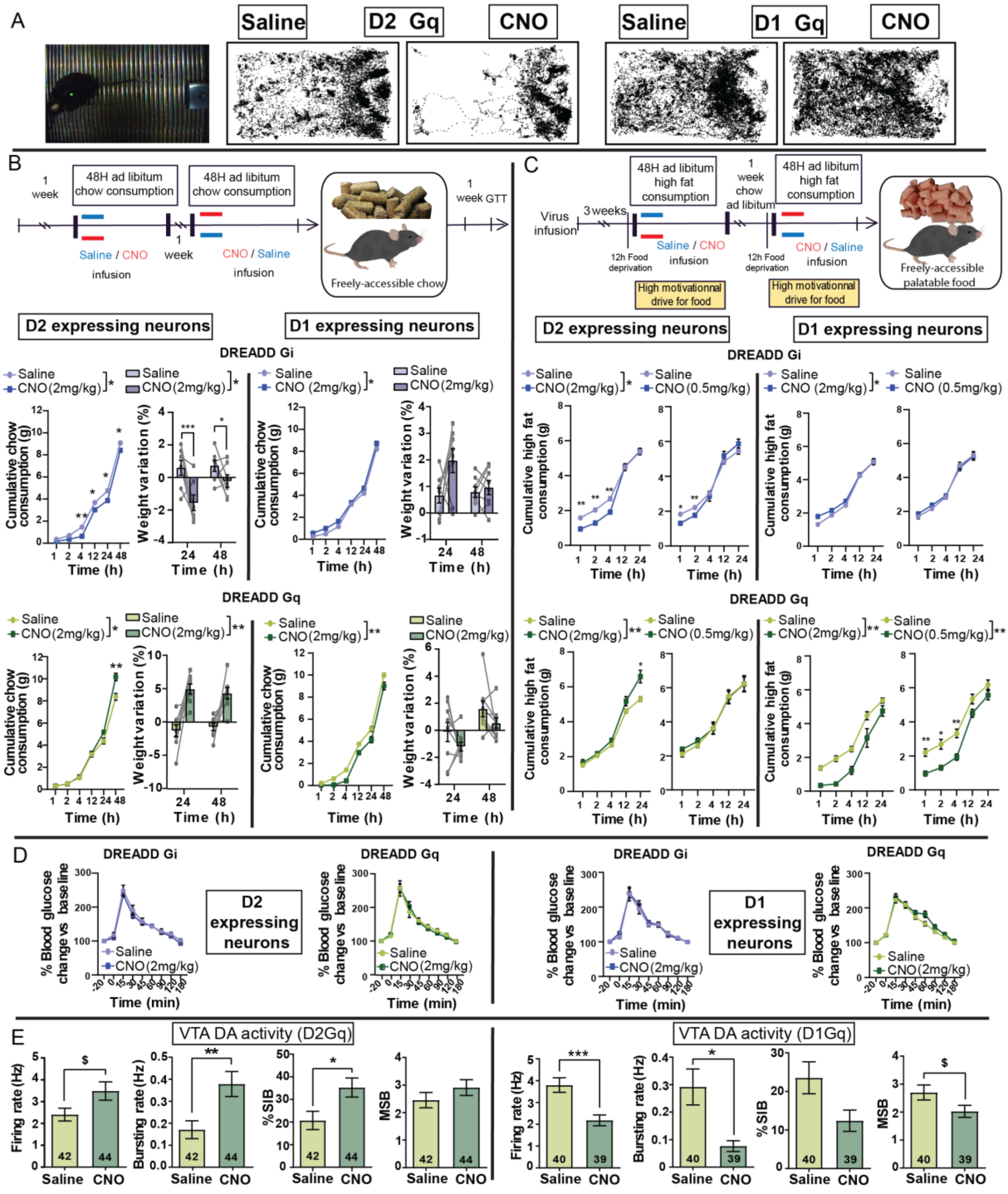


**Suppl. Figure 2 (relates to Figure 2):**

(A) Representative image for the tracking of animal trajectories (left) and examples of trajectories under activation of D2- and D1-neurons comparing saline and CNO conditions from the same animal during pavlovian conditioning. (B) Experimental design (top), and graphs plotting cumulative chow consumption over 48 hours (left column) and weight variations (right column) under inhibtion (DREADD Gi, n=7 for D2-neurons, n=9 for D1-neurons) or activation (DREADD Gq, n=8 for D2-neurons, n=9 for D1-neurons) of D2- or D1-neurons. (C) Fasting-refeeding procedure with high fat diet (top) and graphs plotting cumulative food consumption over 24 hours under inhibtion (DREADD Gi, n=7 for D2-neurons, n=9 for D1-neurons) or activation (DREADD Gq, n=7 for D2-neurons, n=9 for D1-neurons) of D2- or D1-neurons, using 2 doses of CNO. (D) Glucose tolerance test under acute chemogenetic inhibition (DREADD Gi, n=7 for D2-neurons, n=9 for D1-neurons) or activation (DREADD Gq, n=8 for D2-neurons, n=9 for D1-neurons) of NAc D2-neurons (left) and D1-neurons (right). (E) Firing rate, bursting rate, % spike in burst (SIB) and mean spike in burst (MIB) were assessed through In vivo recording of dopaminergic neurons of the VTA under chemogenetic activation of either D2-neurons (n=6) or D1-neurons (n=4) in saline or CNO conditions. Numbers in bars indicate the number of neurons recorded. $: $0.05<p<0.1$ ; * : $p<0.05$ ; ** : $p<0.01$ ; *** : $p<0.001$ ; Error bars = s.e.m. See Table 2 for statistical analyses.

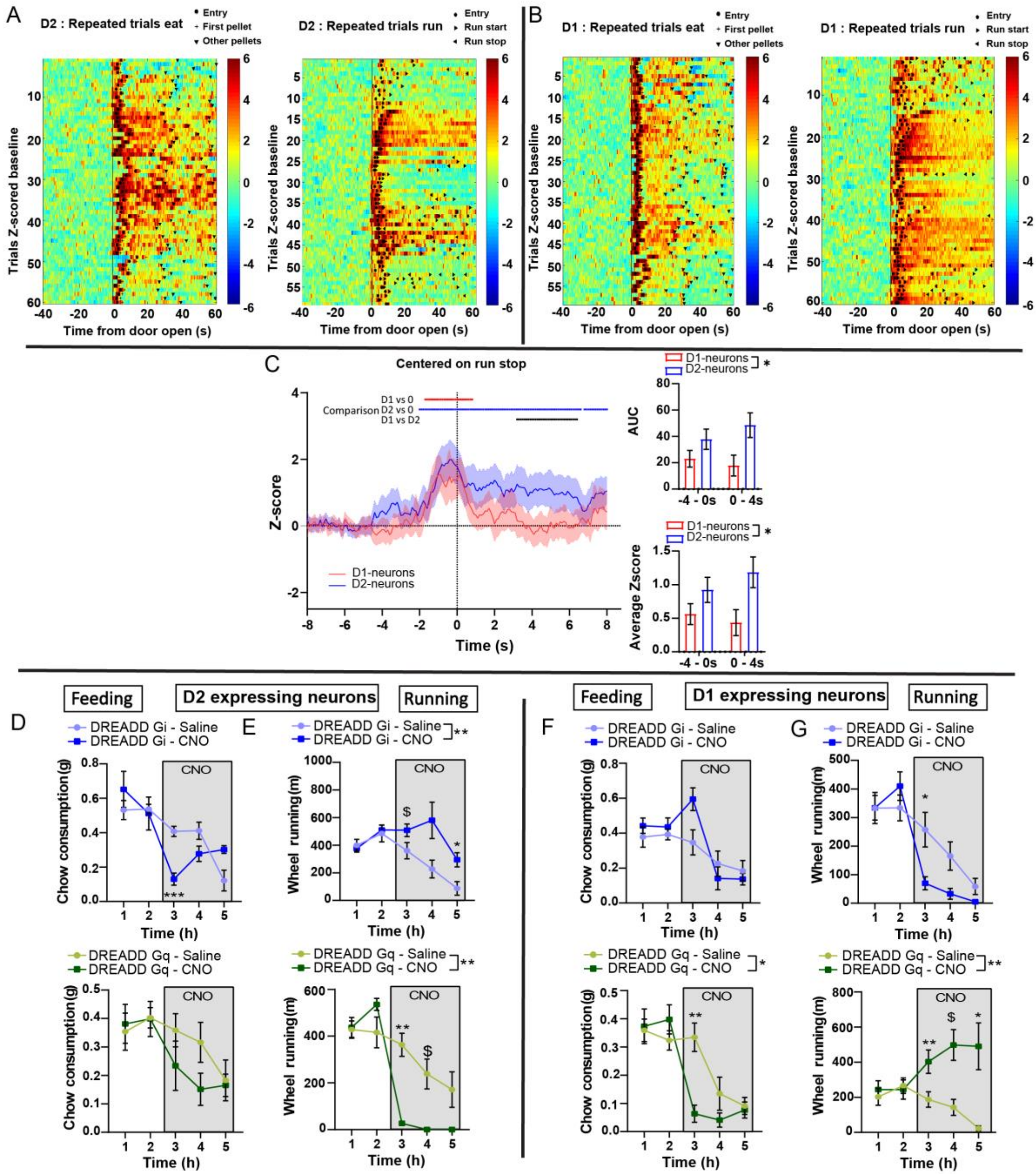


**Suppl. Figure 3 (relates to Figure 3):**

(A) Heat maps of changes in calcium signal (Z score) in D2-neurons (n=5 animals, 12 trials per animal), for feeding (left) or wheel running (right) trials. Events are aligned to door opening. (B) Heat maps of changes in calcium signal (Z score) in D1-neurons (n=5 animals, 12 trials per animal), for feeding (left) or wheel running (right) trials. Events are aligned to door opening. (C) Peri-event analysis of voluntary run stop and analysis of AUC and average z-score. (D-E) Chow consumption (D) and running (E) under chemogenetic inhibition (DREADD Gi, n=7) or activation (DREADD Gq, n=8) of D2-neurons. (F-G) Chow consumption (D) and running (E) under chemogenetic inhibition (DREADD Gi, n=9) or activation (DREADD Gq, n=9) of D1-neurons. * : p<0.05 ; ** : p<0.01; Error bars = s.e.m. See Table 2 for statistical analyses.

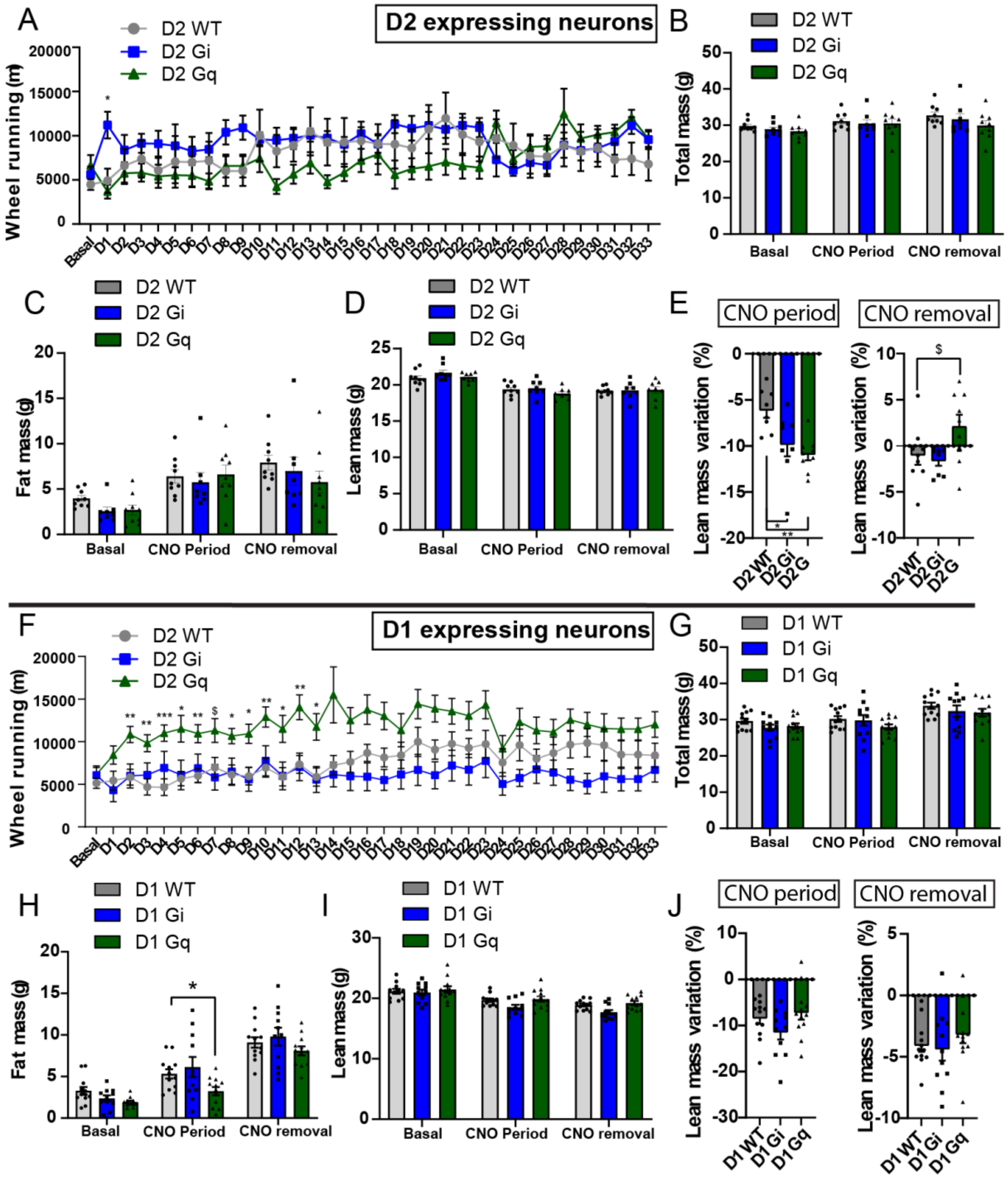


**Suppl. Figure 4 (relates to Figure 4):**

(A-E) Distance ran (A), mass at the end of the CNO period (B), fat mass at the end of the CNO period (C), lean mass at the end of the CNO period (D), and lean mass variations at the end of the CNO period and after CNO removal (E) for chemogenetic manipulations of D2-neurons (WT n=7, D2 Gi n=7, D2 Gq n=8). (F-J) Distance ran (F), mass at the end of the CNO period

(G), fat mass at the end of the CNO period (H), lean mass at the end of the CNO period (I), and lean mass variations at the end of the CNO period and after CNO removal (J) for chemogenetic manipulations of D1-neurons (WT n=9, D1 Gi n=11, D1 Gq n=11). $ : 0.05<p<0.1;  * : p<0.05 ; ** : p<0.01; Error bars = s.e.m. See Table 2 for statistical analyses.

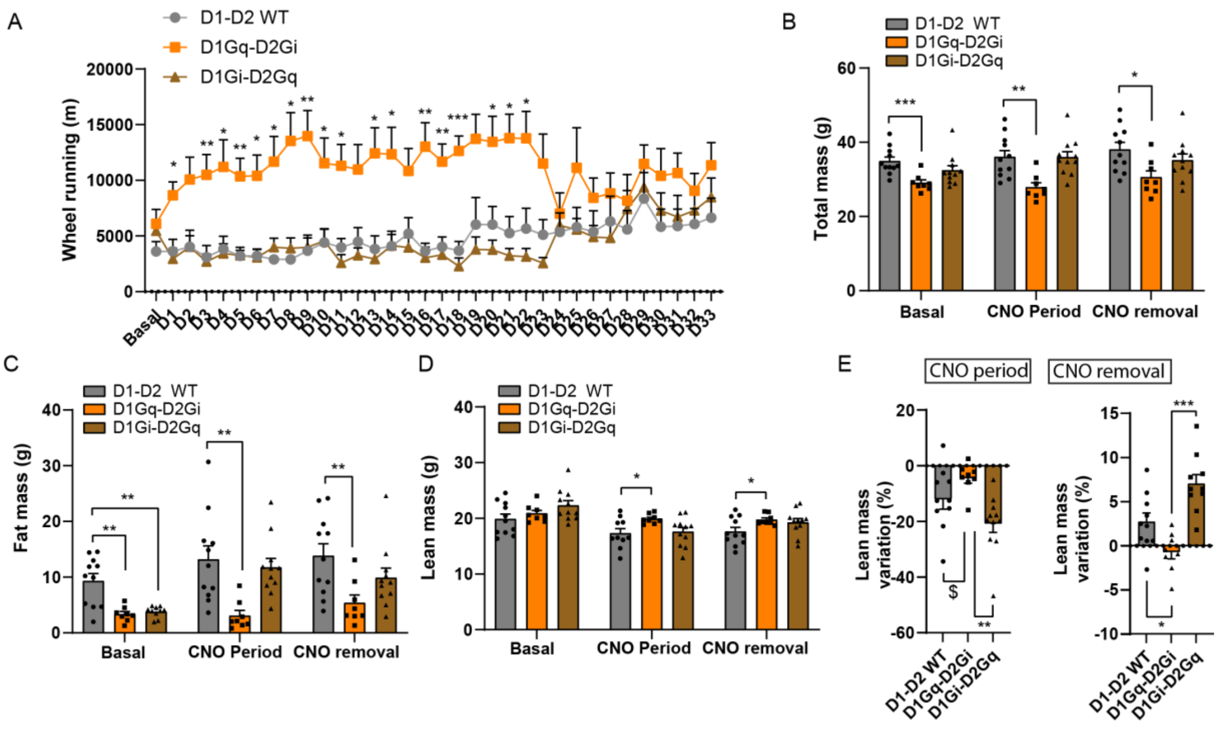


**Suppl. Figure 5 (relates to Figure 5):**

Distance ran (A), mass at the end of the CNO period (B), fat mass at the end of the CNO period (C), lean mass at the end of the CNO period (D), and lean mass variations at the end of the CNO period and after CNO removal (E) for concomitant chemogenetic manipulations of D1- and D2-neurons (WT n=9, D1Gq-D2Gi n=8, D1Gi-D2Gq n=9). $ : 0.05<p<0.1;  * : p<0.05 ; ** : p<0.01; Error bars = s.e.m. See Table 2 for statistical analyses.

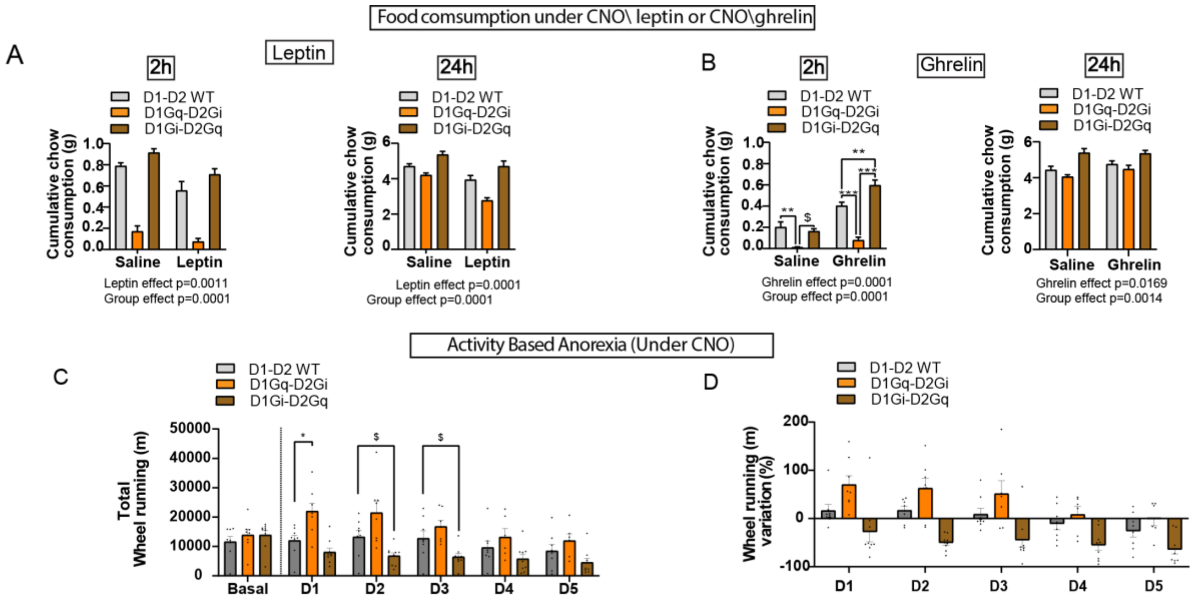


**Suppl. Figure 6 (relates to Figure 6):**

(A) Cumulative chow consumption under leptin administration in saline or CNO conditions under concomitant chemogenetic manipulations of D1- and D2-neurons measured after 2 hours (left) or 24 hours (right) (WT n=8, D1Gq-D2Gi n=7, D1Gi-D2Gq n=7). (B) Cumulative chow consumption under ghrelin administration in saline or CNO conditions under concomitant chemogenetic manipulations of D1- and D2-neurons measured after 2 hours (left) or 24 hours (right) (WT n=8, D1Gq-D2Gi n=7, D1Gi-D2Gq n=7). (C-D) total distance ran (C) and wheel running variation (D) over the ABA procedure under concomitant chemogenetic manipulations of D1- and D2-neurons (WT n=8, D1Gq-D2Gi n=8, D1Gi-D2Gq n=9). $ : 0.05<p<0.1; * : p<0.05 ; ** : p<0.01; Error bars = s.e.m. See Table 2 for statistical analyses.
